# Effects of breed, management and personality on cortisol reactivity in sport horses

**DOI:** 10.1101/739847

**Authors:** Fay J. Sauer, Marco Hermann, Alessandra Ramseyer, Dominik Burger, Stefanie Riemer, Vinzenz Gerber

## Abstract

Sport horses need to fulfill high physical and psychological requirements during training and competition. These as well as certain conditions of modern husbandry may affect their wellbeing. Here we aimed to (1) evaluate effects of demographic and management factors as well as personality traits on stress reactivity of sport horses, (2) investigate if elite sport horses have elevated stress levels compared to amateur sport horses, and (3) assess whether different equestrian disciplines differentially influence horses’ adrenal cortex responsiveness. For this purpose, we visited 149 healthy elite (*n*=94) and amateur (*n*=54) sport horses in Switzerland and performed an adrenocorticotropic hormone (ACTH) stimulation test. Additionally, a person who was familiar with the horse completed a questionnaire about demographic and management factors and horses’ personality traits. Linear models were calculated to assess associations between the questionnaire data and salivary cortisol 60 and 90 minutes after ACTH stimulation. While the model at T90 was not significant, post-stimulatory cortisol after 60 min appears most informative in line with a previous study and was significantly affected by the horses’ breed and by three management factors: “number of riders”, “hours spent outside” and “group housing” (adjusted r^2^ =15%, p<0.001). Thoroughbred and Warmblood horses displayed an increased adrenal response compared to Franches-Montagnes horses. Horses with several riders had a less pronounced reaction than horses with one rider, and horses that spent more time outside had a decreased response compared to horses that were stabled most of the time. Horses living in groups showed higher post-stimulatory cortisol values than horses that were housed singly. However, no significant associations of cortisol responsiveness with personality traits were found, and neither the horses’ use as elite or as amateur sport horses nor the discipline had an effect on the cortisol response. This suggests that optimizing husbandry conditions may be more important for improving horses’ welfare than changing their use.

## Introduction

As grazing animals living in herds [1, 2], their physical strength, speed and endurance enabled horses (*Equus caballus*) to flee from predators and to search for food and water over large distances in their original steppe environment [3]. Within human custody, these naturally occurring talents predestined horses to become extraordinary athletes [3]. Furthermore, these features have even been enhanced through selective breeding for competition purposes and training [4, 5]. However, modern keeping conditions often bear little resemblance to the horse’s natural habitat and social structure and in many cases allow only limited natural foraging behavior [1]. Sport horses, especially, are now mostly housed in single stalls [6] and their time on pasture and the amount of roughage fed is often restricted [7]. In equestrian sports, horses often complete high intensity trainings with different trainers and riders, go to national and international horse shows, and are thus often required to live in different stables and to travel long distances in trailers and airplanes. As a consequence, modern sport horses are significantly challenged [8]. The high requirements that they are expected to fulfill, as well as the conditions of modern husbandry, may lead to significant stress, both acute and chronic.

Stress is the response of the body to a demand (stressor) placed upon it [9]. Whenever a stimulus is perceived as a stressor, it results in a biological response in an attempt to cope with the situation [10]. This biological reaction consists of the behavioral response, the sympathetic-adrenal medulla (SAM) axis response and the hypothalamic-pituitary-adrenal cortex (HPA) axis response, which are often activated concurrently and interact during stressful situations [11]. Studies investigating acute or chronic stress in horses have assessed either one or several of these pathways [12–14]. SAM axis parameters, such as evaluation of heart rate or heart rate variability and endocrinological parameters such as cortisol levels are typically used as measures of acute stress [10]. HPA axis stimulation tests such as adrenocorticotropic hormone (ACTH) or corticotropin releasing factor (CRF) challenge tests are described as potential tools for the assessment of long-lasting effects of stress in horses [10]. Stimulated cortisol may give an impression of an animal’s overall adrenal capacity, with both a depression [15–17] or an exaggerated response to stimulation [18–21] reflecting a long-term stress-related alteration of the HPA axis.

Several management factors as well as personality traits have been investigated in relation to stress responsivity in horses. For instance, a lack of social contact was linked to changes in stress-related physiology and behavior [22, 23]. The examination of feeding patterns revealed associations between stereotypies as indicators of chronic stress [10] and a restriction of forage intake [24, 25] or pasture time [26–28]. Furthermore, participation in equestrian competitions was demonstrated to cause an increased sympathoadrenal activity and an activation of the HPA axis immediately after the event [29–31]. A few studies also reported some associations between equine personality traits and stress parameters [32–34]. Cortisol responses following external stimulation of the HPA axis have been investigated in behavioral studies [23, 35] and for the evaluation of chronic stress in horses suffering from overtraining [36], stereotypies [18] or gastric disease [19, 20].

However, to what extent management factors, type of use or horses’ personality traits are associated with indicators of long-term stress has not been investigated to date. Using ACTH stimulation tests and questionnaire surveys, the aims of this study were to explore (1) effects of breed, management factors and personality traits on the adrenal reactivity of sport horses, (2) whether elite sport horses have elevated stress-levels in comparison to amateur horses, and (3) if sport discipline is an influencing factor.

## Materials and methods

### Sample population

The study was approved by the cantonal authority for animal experimentation, the Veterinary Office of the Canton of Bern, Switzerland, and subsequently by all other included cantons (27608/BE26/16). Before participating in the study, all owners gave their informed written consent. This prospective cross-sectional study was carried out between January and August 2017. Swiss private practitioners and official team veterinarians of the different disciplines were contacted to establish contacts with riders and owners in order to recruit horses of different equestrian disciplines (show jumping, dressage, eventing, endurance, driving, vaulting, para-equestrian) for both an Elite Sport Horse Group (ESHG) and an Amateur Sport Horse Group (ASHG), respectively. In order to participate in the study, horses had to have competed at predefined levels of competition (Table 1). Overall, 149 horses from 17 different cantons in Switzerland were included. Ninety-five of them were part of the ESHG and 54 of the ASHG (Table 1).

**Table 1.** Horses included in the study and inclusion criteria.

| Discipline | Number of horses | Breeds |  |  |  | Required competition level of horses |
| --- | --- | --- | --- | --- | --- | --- |
| W F T O |  |  |  |  |  |  |
| Sport Horse Group: |  |  |  |  |  |  |
| Show Jumping | 17 | 16 |  | 1 |  | Concours de Saut Internationaux (CSI) * to ***** |
| Dressage | 14 | 13 |  |  | 1 | Concours de Dressage Nationaux (CDN) and Concours de Dressage Internationaux (CDI)* to ***** |
| Eventing | 20 | 17 |  | 3 |  | Concours Complet Nationaux (CCN) and Concours Complet Internationaux (CCI) * to *** |
| Endurance | 20 |  |  | 20 |  | Concours d’Endurance Internationaux (CEI) * to *** |
| Driving | 14 | 4 | 10 |  |  | FEI approved international class M and S |
| Vaulting | 5 | 5 |  |  |  | Concours de Vaulting Internationaux (CVI) * to ***** |
| Para Equestrian | 5 | 5 |  |  |  | FEI-approved international class M and S |
| Amateur Horse Group: |  |  |  |  |  |  |
| Show Jumping | 16 | 16 |  |  |  | National competition level 90 to 120 cm |
| Dressage | 18 | 15 | 2 |  |  | National competition level GA03/05 to L |
| Eventing | 9 | 9 |  |  |  | National competition level B1 to B3 |
| Endurance | 11 |  |  | 10 | 1 | National competition level EVG1 to EVG4 |

List of horses of the Elite Sport Horse Group (ESHG) and the Amateur Sport Horse Group (ASHG) that participated in the study. For every discipline, the number of animals and the breeds of the participating horses is included and the required level of competition (inclusion criteria) is given. W = Warmblood horses; F = Franches-Montagnes horses, T = Thoroughbred horses; O = Others.

### Experimental setup

The first author visited all horses in their “home stable” environment. Each horse underwent an ACTH stimulation test, and a person who was familiar with the horse (owner, rider or caretaker) completed a questionnaire. Horses did not participate in any physical activity two hours before the visit, and they were not transported and did not attend competitions during the preceding 48 hours. They received no medication or additives except for over-the-counter nutritional supplements. Prior to the ACTH stimulation test, the weight of the horses was estimated using an established formula for mature horses, where weight (kg) = (*heartgirth^2^ x body length*)/ (*11,880 cm*^3^) [37]. The general examination included an assessment of mucous membrane color, lymph nodes, heart rate, respiratory rate and rectal temperature at rest. Only horses in a good general condition with normal general examination findings were included in the study. The body condition score (BCS) was recorded, according to the 9-point Henneke BCS system ranging from 1 (poor) to 9 (extremely fat) [38]. During the time of the test, horses remained in their normal environment such as their stall or open stable. They were prevented from drinking during the time of the test to avoid dilution of saliva before cortisol measurements. For this reason, self-watering troughs were covered with plastic bags.

### ACTH stimulation test

For the ACTH stimulation test, based on the estimated weight, a dose of 1 μg/kg BW synthetic ACTH1-24 (Synacthen tetracosactidum 0.25 mg/ml equivalent to 25 IU/mL; Novartis, Vilvoorde, Belgium) was injected intravenously. The dosage was chosen based on previous publications [19, 20, 39, 40]. Saliva samples were taken with salivettes (Sarstedt, Nuembrecht, Germany) before (0 minute – baseline; T0) and 60 (T60) and 90 (T90) minutes after ACTH administration. The salivette swab was placed into the horse’s mouth for at least 40 seconds until it was completely soaked with saliva. Afterwards, the swab was replaced into the salivette and stored in a box with cool packs until return to the clinic. Salivettes were then directly centrifuged at room temperature for 10 min at 185 x *g* and stored at −20°C until the analysis. For the determination of salivary cortisol concentrations, a competitive enzyme immunoassay (cELISA, Salimetrics, Newmarket, United Kingdom) was used, which has been previously validated for use in horses [19, 20, 40].

### Questionnaire

The first part of the questionnaire was designed to record information about demographic and management factors such as housing, use and feeding, stereotypies and clinical signs such as teeth grinding or yawning (Table 2). After termination of the study, some questions were re-coded because there were too few observations in one of the categories (Table 2). The aim of the second part of the questionnaire was to evaluate personality traits of the participating horses. For this purpose, a previously validated questionnaire [41], relying on the opinion of caretakers regarding the personality of horses, was adapted for the current study (Table 3). The original version by Momozawa et al. [41] consisted of 20 items, leading to four factors after a principal component analysis. These factors were labelled ‘Anxiety’, ‘Trainability’, ‘Affability’ and ‘Gate Entrance’. For the current study, the question about the horses’ behavior at the starting gate (“gate entrance”) was excluded, since it is only relevant for racehorses. Three additional questions regarding cooperation during riding, trailer loading and transportation and four questions about the horses’ resilience during or after putatively stressful events such as competitions or transports including [1] recovery time; [2] susceptibility to infection (e.g. respiratory infections); [3] loss of appetite; and [4] signs of discomfort (e.g. flehmen, colic, prolonged resting times) were added instead. Questions of this second part of the questionnaire were answered using a scale ranging from (1) poor to (9) excellent. To identify possible pitfalls or misunderstandings, four independent volunteers/horse owners completed both parts of the questionnaire. The questionnaire was adapted according to their comments, and German and French translations were prepared for the participants. A person who was familiar with the horse was asked to complete the questionnaires during the time the ACTH stimulation test was performed.

**Table 2.**
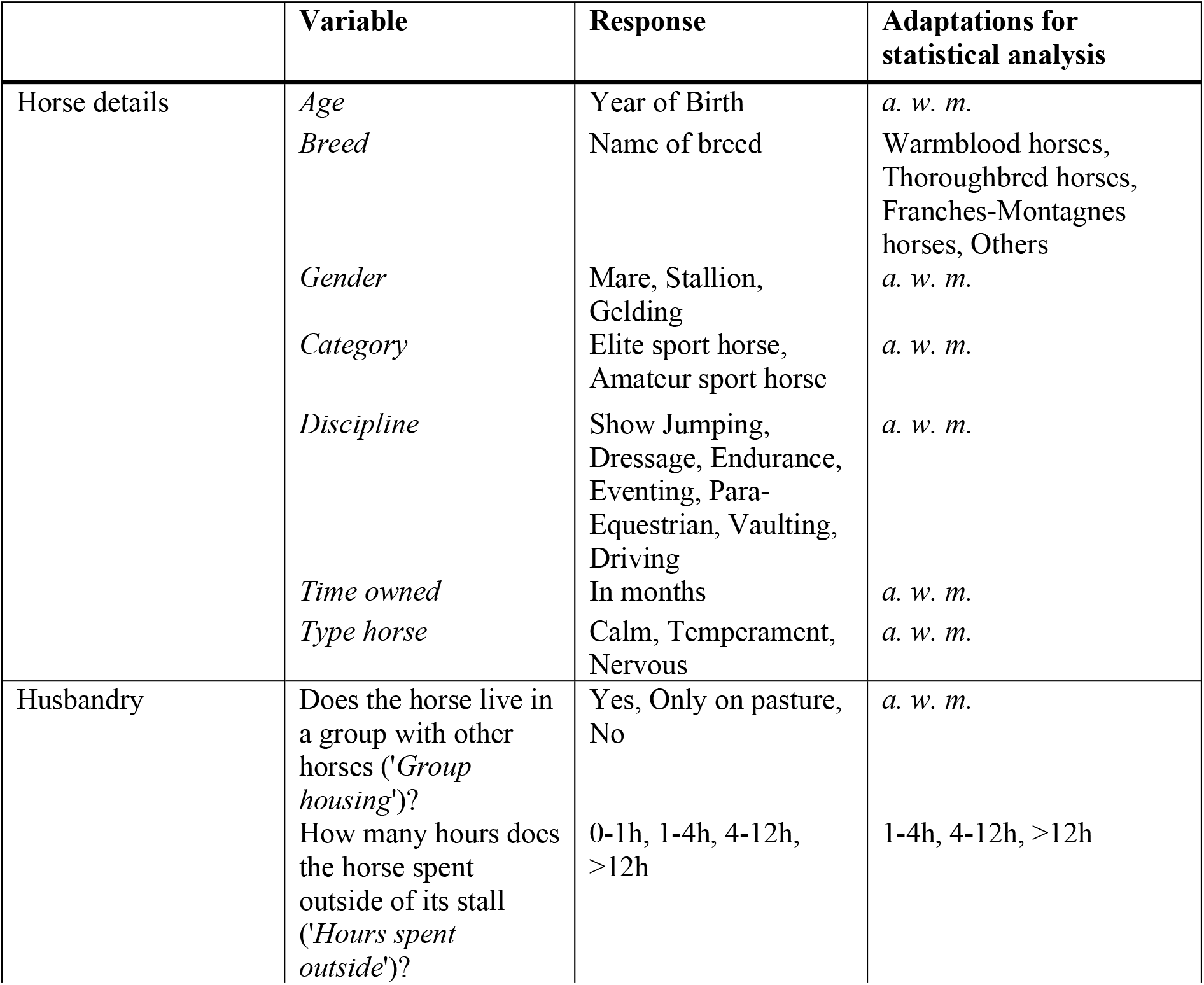
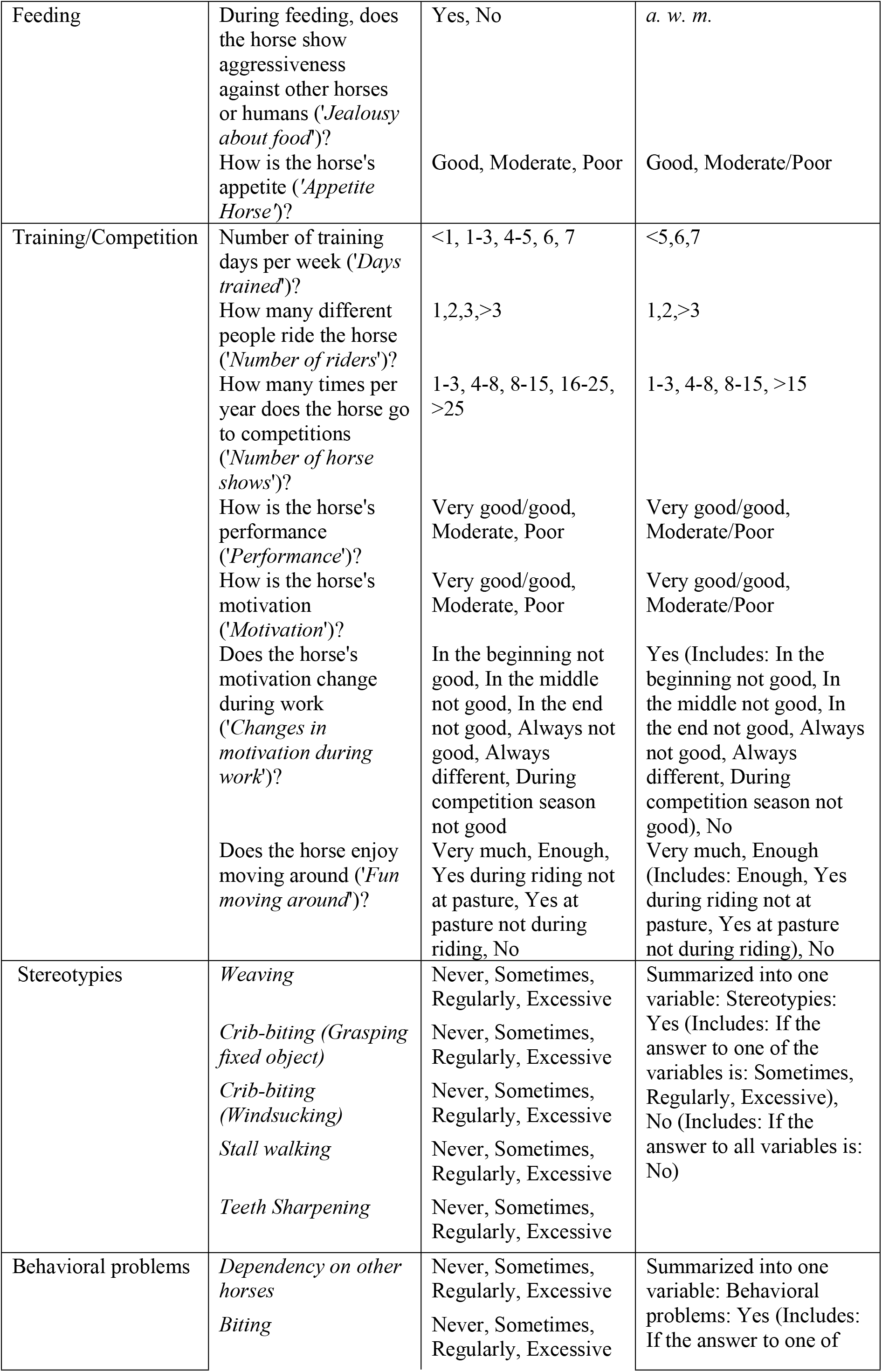
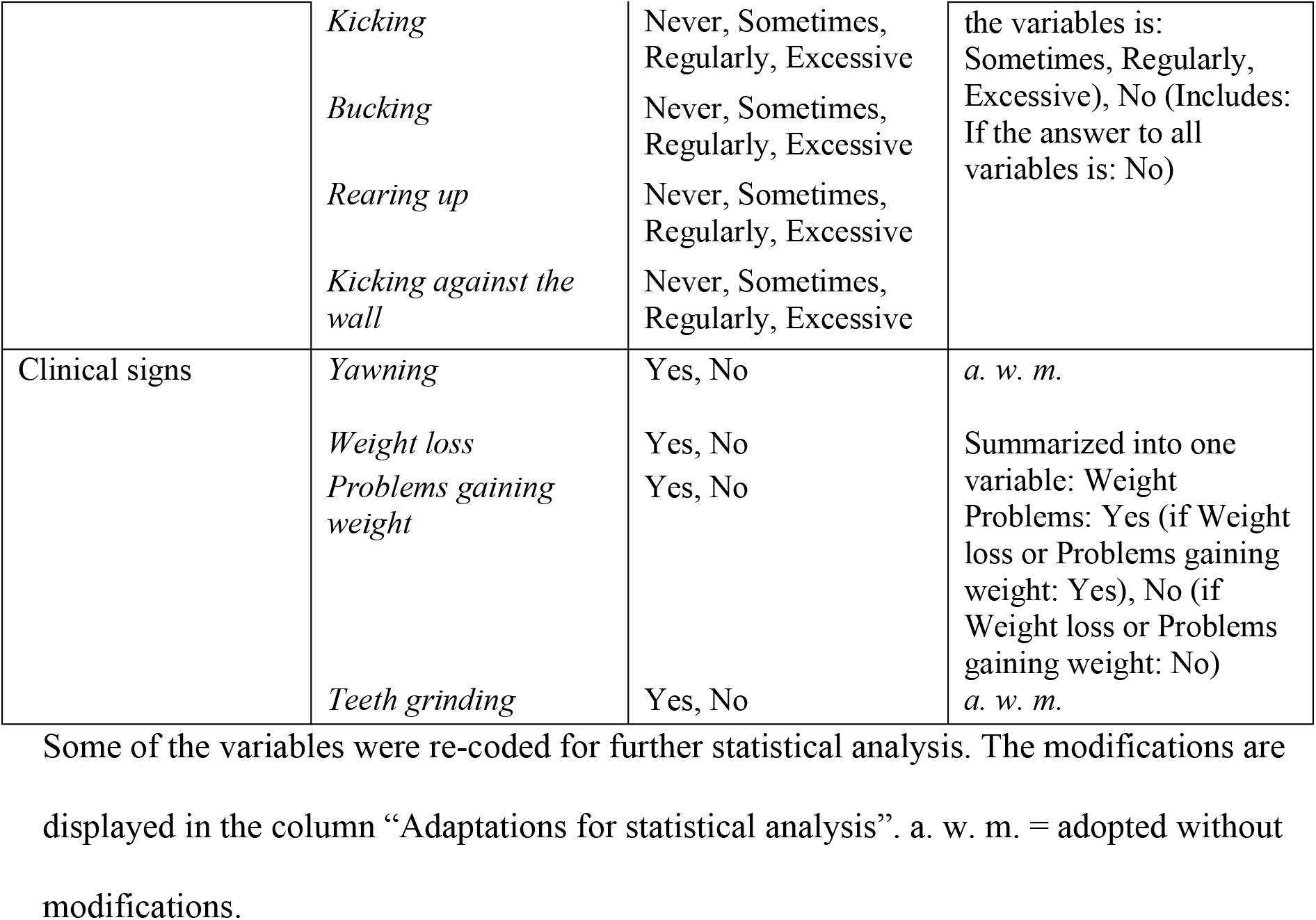
First part of the questionnaire containing questions about demographic and management factors.

### Statistical analysis

All statistical analyses were performed in R [42].

### Principle Component Analysis on personality traits

A Principle Component Analysis PCA (function prcomp) was performed on the second part of the questionnaire (personality traits) in order to reduce the number of variables and obtain principle components for further analysis. A Varimax rotation was carried out, and components with Eigenvalues >1 were retained.

### Evaluation of the effects of demographic and management factors as well as personality traits on stimulated cortisol levels

Linear regression models (function lm) were calculated separately for the dependent variables T60 and T90, with separate models for (1) demographic and management factors and (2) personality traits, respectively. In the first model, 24 demographic and management factors from the first part of the questionnaire (Table 2) in addition to the exact time of day of the start of the ACTH stimulation test and the horse’s BCS were included as predictors. In the second model, the seven principal components derived from the personality questionnaire were used as explanatory variables. Initially, all variables were entered in the models, which were then reduced by stepwise backward selection (function step). The fit of the models was assessed using Akaike’s Information Criterion (AIC), and the model with the lowest AIC was retained as the final model. Level of significance was set at p < 0.05. Results were corrected for multiple testing via Bonferroni correction. In the Results section, we report original p-values and subsequently indicate whether they can still be considered significant after Bonferroni correction. The residuals of the models were checked visually via scatterplots for independence and via quantile-quantile-plots, histograms and Shapiro-Wilk tests for normal distribution. The models were tested for homoscedasticity of variance via visual inspection and Breusch-Pagan test (package car, function ncvTest). If necessary, square-root transformation of the respective response variable was performed to meet linear model assumptions. Variables of the models were checked for multicollinearity via use of the variance inflation factor (package car, function vif). There was no multicollinearity in the variables included in the final models. 95% confidence intervals and likelihood-ratio tests (LRT) were calculated for the final reduced models.

### Evaluation of the effect of competition level and discipline on stimulated cortisol levels

For the comparison of adrenal reactivity in the ESHG versus the ASHG, the disciplines para-equestrian, driving and vaulting were excluded, since no comparable groups of amateur sport horses were available. Thus, 71 horses of the ESHG and all 54 horses of the ASHG were included in the analysis. Linear regression models were computed to evaluate the effect of category (ESHG versus ASHG) and the influence of the different disciplines on cortisol levels after exogenous stimulation. As above, model assumptions were checked and were adequate for all models.

## Results

### Sample population

Data are reported as mean ± standard deviation (range). Horses in the ESHG were 11 ± 3 (6 to 22) years old and consisted of Warmbloods (n = 60), Thoroughbreds (n = 24), Franches-Montagnes horses (n = 10) and one horse of another breed. Fifty-seven were geldings, 33 mares and 5 stallions. Horses of the ASHG were 11 ± 3 (5 to 14) years old and included the breeds Warmblood (n = 40), Thoroughbred (n = 10), Franches-Montagnes horses (n = 2), and two horses of other breeds (Table 1). The sample included 28 geldings, 24 mares and 2 stallions.

### Principal Component Analysis on personality traits

Out of the 26 questions of the second part of the questionnaire, those on ‘competitiveness’ and ‘docility’ were removed. Many horses did not have direct contact with other horses; therefore, it was not possible to evaluate competitiveness. German and French translations of docility turned out to be unsatisfactory and respondents were confused about the meaning. The PCA on the remaining 24 items yielded seven cumulative factors with an Eigenvalue > 1 accounting for 63.4% of the common variance. The factors partially confirmed the factor structure determined for the original version of the questionnaire [41]. Factor 1 included five out of six items (nervousness, excitability, panic, skittishness, timidity) that were grouped as “anxiety” by Momozawa et al. [41]. In the present study, it contained two additional items (concentration, perseverance). Factor 2 (“trainability”), consisted of two (trainability, memory) out of four items grouped by Momozawa et al. and of two additional items (stubbornness, cooperation during riding). Factor 3 consisted of four items that were not part of the original questionnaire and was named “recovery” (recovery time after efforts, infections after efforts, inappetence after efforts and discomfort after efforts). Factor 4, “social interaction” and factor 5, “alertness”, did not form part of the factors identified by Momozawa et al. Factor 4 consisted of “self-reliance”, “friendliness towards horses” and “discomfort after efforts” and factor 5 of “concentration”, “curiosity” and “vigilance”. In agreement with Momozawa et al., Factor 6 was labeled “affability”. It consisted of two (friendliness towards people, cooperation while taking care) out of the four items that were grouped in the original questionnaire and of one further item (inconsistent emotionality). The newly defined factor 7, “transportability”, consisted of the newly added questions “cooperation trailer loading” and “cooperation transport” (Table 3 and Fig 1).

**Table 3.**
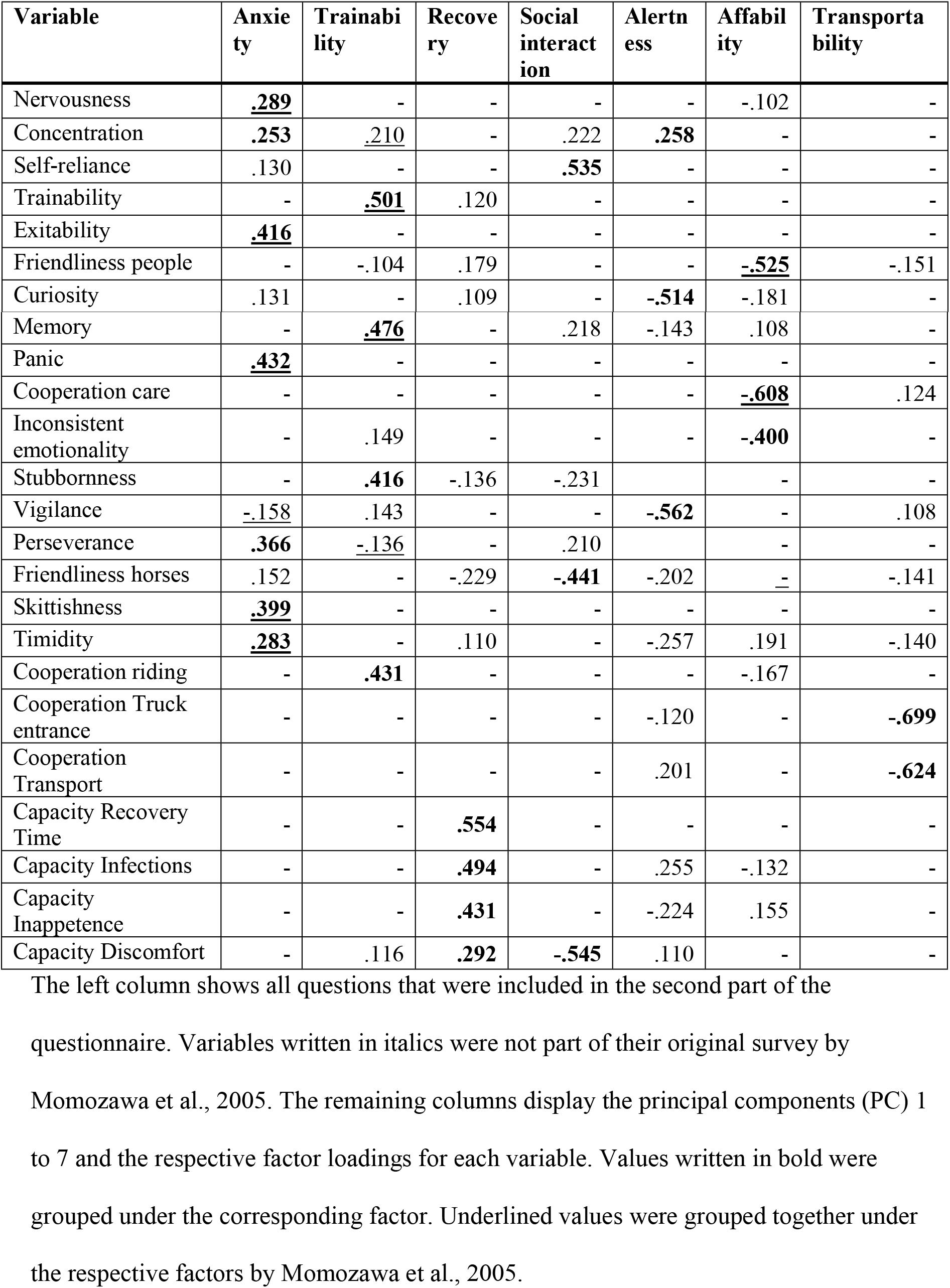
Factor loadings of the Principal Component Analysis on personality factors.

**Fig 1.**
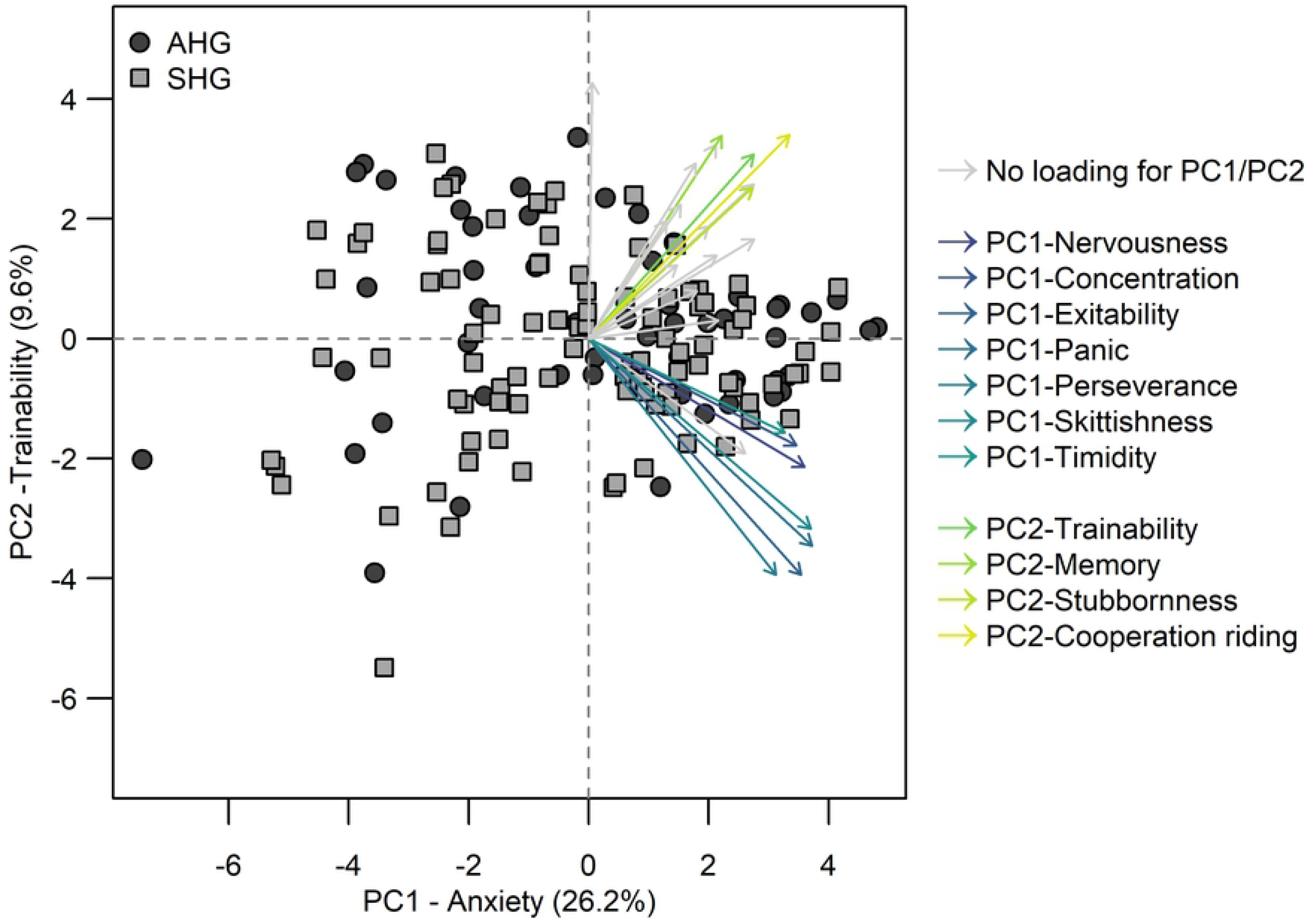
Principal component 1 (PC1-Anxiety) versus 2 (PC2-Trainability) of the Principal Component Analysis on personality traits. Horses of the Elite Sport Horse Group (ESHG) are depicted as dark grey dots and horses of the Amateur Sport Horse Group (ASHG) as light grey squares. Arrows of the variables accounting for PC1-Anxiety are visualized in shades of purple and blue and arrows of the variables accounting for PC2-Trainability are shown in yellow to green tones. Arrows of variables that did not contribute to PC1-Anxiety or PC2-Trainability are grey.

### Evaluation of the effects of demographic and management factors as well as personality traits on stimulated cortisol levels

The final reduced linear regression model evaluating the effect of demographic and management factors on cortisol levels 60 minutes after ACTH stimulation was significant after Bonferroni correction (T60: p < 0.001; Adjusted R^2^ = 15%; F_1, 122_ = 3.305). Likelihood ratio tests revealed four variables with a significant effect: ‘breed’ (LRT: p = 0.02), ‘number of riders’ (LRT: p = 0.009), ‘hours spent outside’ (LRT: p = 0.002) and ‘group housing’ (LRT: p = 0.045) (Table 4). Regarding ‘breed’, positive parameter estimates for Warmblood and Thoroughbred horses indicate higher post-stimulatory cortisol values in these breeds compared to Franches-Montages horses that served as the reference in the model. Regarding the ‘number of riders’, horses with one rider displayed higher post-stimulation cortisol values than horses with two or more riders. Furthermore, horses that spent more time outside of their stalls on pasture or for training showed lower values than horses being less outside. Predicted values for cortisol 60 minutes after ACTH stimulation are plotted for the predictors ‘breed’, ‘number of riders’, ‘hours spent outside’, and ‘group housing’ in Fig 2.

The linear regression model evaluating demographic and management factors influencing cortisol levels 90 minutes after ACTH stimulation (Table 4) was not significant (T90: p = 0.085; Adjusted R^2^ = 4%; F_1, 122_ = 1.897).

When considering the correction for multiple testing, none of the personality traits was significantly associated with post-stimulatory cortisol at either 60 or 90 minutes (T60: p = 0. 043, adjusted R^2^ = 2%; F_1, 122_ = 4.181; T90: p = 0.013, adjusted R^2^ = 4%, F_1, 122_ = 6.35)) (Table 5).

**Table 4.**
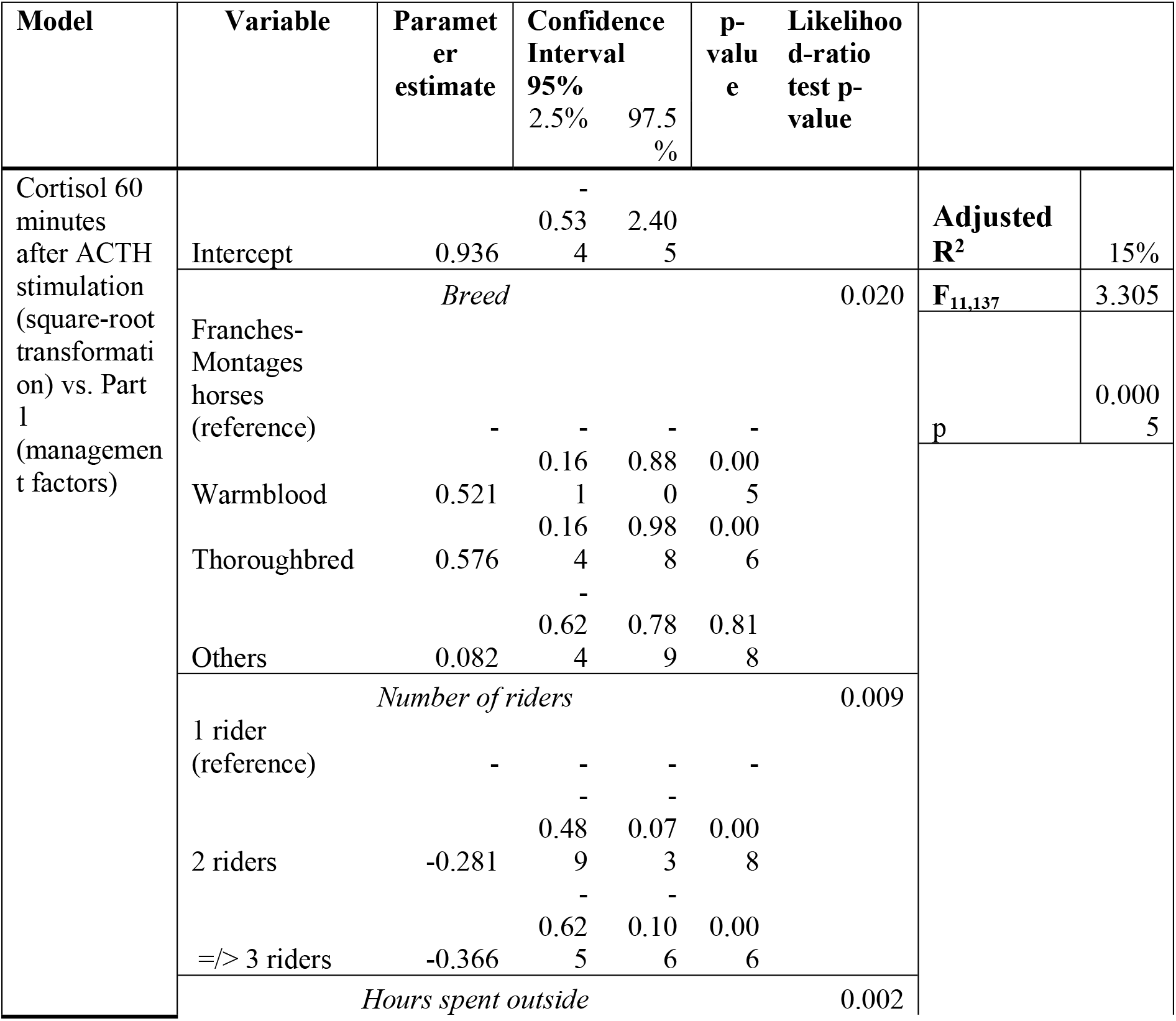
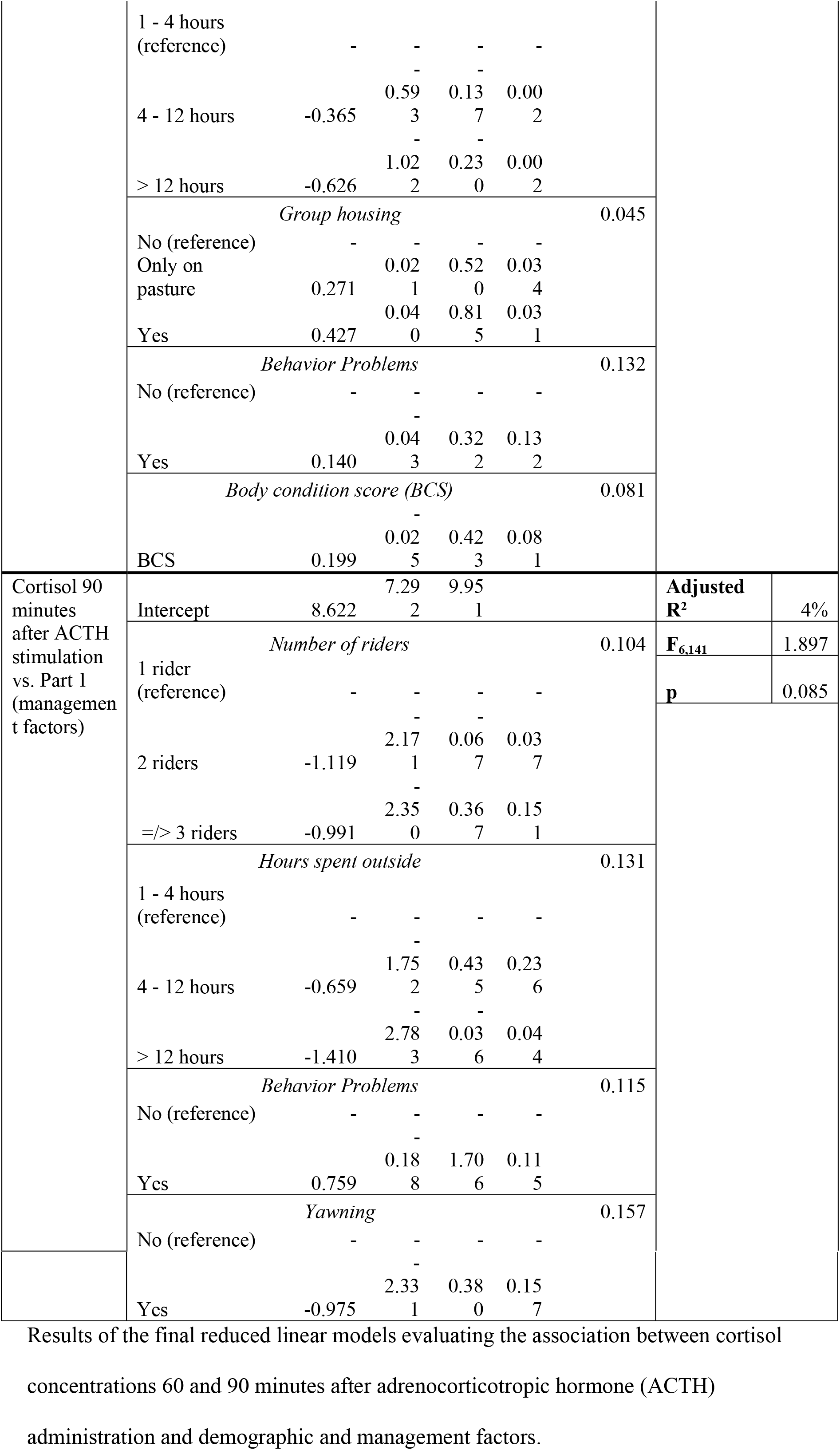
Effects of demographic and management factors on cortisol concentrations after ACTH stimulation.

**Fig 2.**
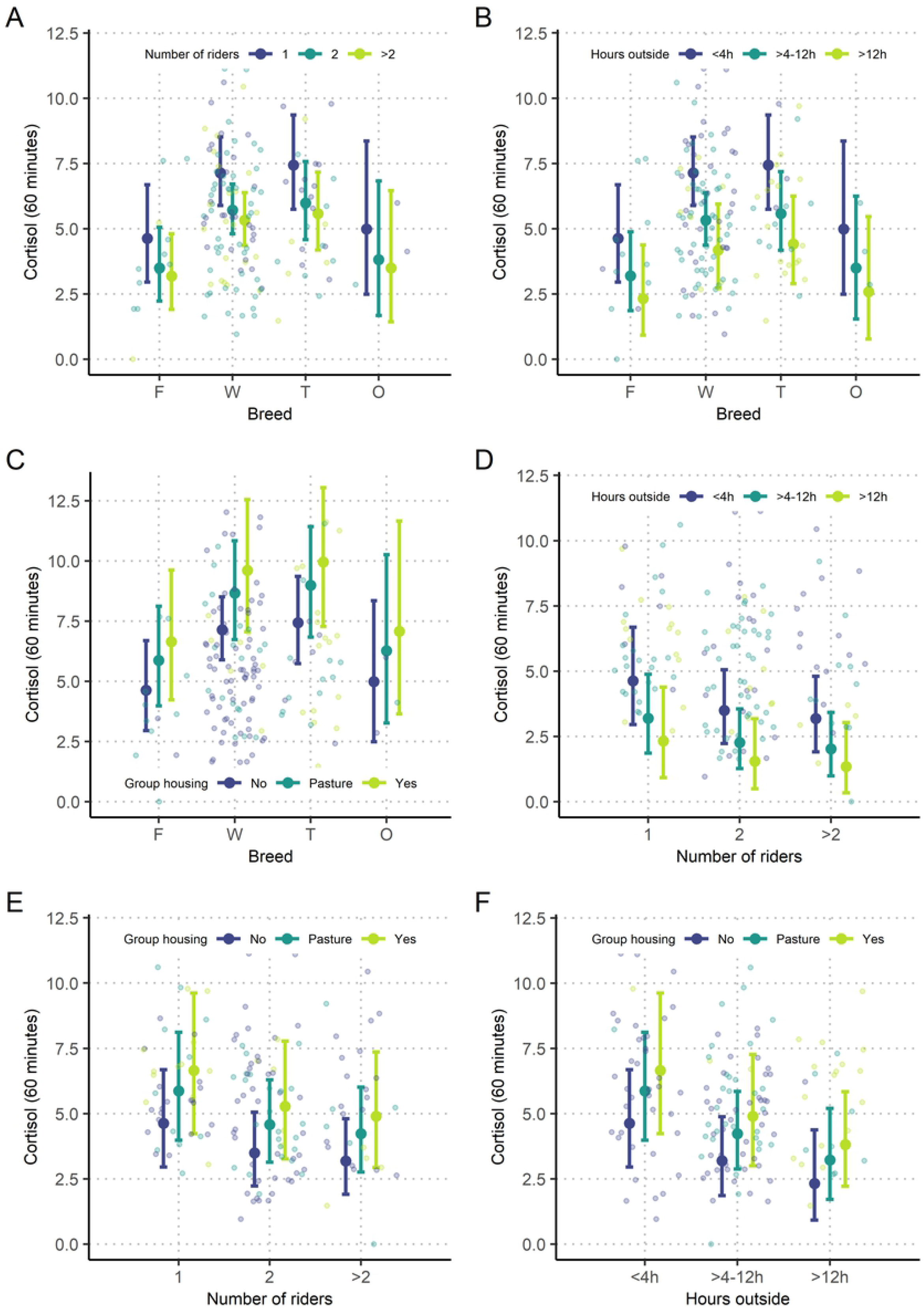
Predicted values of cortisol based on the linear regression model at 60 minutes after ACTH stimulation. The explanatory categorical variables ‘breed’, ‘number of riders’, ‘hours spent outside’ and ‘group housing’ are plotted against each other for prediction of cortisol values, 60 minutes after ACTH stimulation: The six plots display predicted values of cortisol for (A) ‘breed’ versus ‘number of riders’, (B) ‘breed’ versus ‘hours spent outside’, (C) ‘breed’ versus ‘group housing’, (D) ‘number of riders’ versus ‘hours spent outside’, (E) ‘number of riders’ versus ‘group housing’ and (F) ‘hours sent outside’ versus ‘group housing’. Open dots indicate underlying raw data.

**Table 5.** Final reduced models assessing effects of personality factors on cortisol concentrations after ACTH stimulation.

| Model | Variable | Parameter estimate | Confidence Interval 95% |  | p-value |  |  |
| --- | --- | --- | --- | --- | --- | --- | --- |
|  |  |  | 2.5% | 97.5% |  |  |  |
| Cortisol 60 minutes after ACTH stimulation (square-root transformation) vs. Part 2 (personality factors) | Intercept | 2.297 | 2.206 | 2.389 |  | Adjusted R<sup>2</sup> | 2% |
|  | *Social interaction* |  |  |  |  | F<sub>1,147</sub> | 4.181 |
|  | Social interaction | -0.079 | -0.155 | -0.003 | 0.043 | p | 0.043 |
| Cortisol 90 minutes after ACTH stimulation vs. Part 2 (personality factors) | Intercept | 7.390 | 6.947 | 7.833 |  | Adjusted R<sup>2</sup> | 4% |
|  | *Social interaction* |  |  |  |  | F<sub>1,146</sub> | 6.350 |
|  | Social interaction | -0.471 | -0.841 | -0.102 | 0.013 | p | 0.013 |

Results of the final reduced linear models evaluating the association between cortisol concentrations 60 and 90 minutes after adrenocorticotrophic hormone (ACTH) administration and the factors anxiety, trainability, recovery, social interaction, alertness, affability and transportability gained by Principal component analysis of the second part of the questionnaire about personality traits.

### Evaluation of the effect of competition level and discipline on stimulated cortisol levels

There was no significant effect of the category (ESHG or ASHG) on cortisol values after ACTH stimulation (T60: p = 0.819; adjusted R^2^ = −0.008; F_1, 123_ = 0.05 and T90: p = 0.721; adjusted R^2^ = −0.007; F_1, 123_ = 0.13). Likewise, cortisol levels at T60 (p = 0.753; adjusted R^2^ = −0.02; F_7, 117_ = 0.6) and T90 (p = 0.055; adjusted R^2^ = 0.06; F_7, 117_ = 2.05) did not differ between horses of different disciplines.

## Discussion

This study found that ‘breed’, ‘number of riders’, ‘hours spent outside’ and ‘group housing’ are significantly related to salivary cortisol concentrations in horses 60 minutes after ACTH stimulation, which is considered as a measure of long-term effects of stress [10]. There was, however, no difference in post-stimulatory cortisol levels between elite sport horses and amateur sport horses, nor between horses of different disciplines. Furthermore, personality traits were unrelated to horses’ cortisol responsiveness.

Notably, 60 minutes post-stimulation appeared to be the most informative time point, as linear regression models evaluating cortisol concentrations after 90 minutes showed no significant outcome. This is in agreement with our previous study [43], where sampling after 60 minutes also revealed the best association with the occurrence of glandular gastric disease. The present study supports our previous hypothesis [43] that predominantly the initial increase of cortisol after ACTH stimulation is important for the evaluation of the cortisol responsiveness in the context of stress-related disorders.

Thoroughbreds and Warmbloods displayed an increased adrenal response in comparison to Franches-Montagnes horses. This is in line with a previous study of our group, finding a lower response to ACTH in Franches-Montagnes horses compared to Warmblood horses [40]. In this previous study, however, all Franches-Montagnes horses were stallions and so it was not possible to differentiate whether the effect was due to the sex or the breed of the horses. The present study investigated the potential effects of sex and breed as separate explanatory variables, indicating that the breed (but not sex) might be influencing equine adrenal reactivity. This suggests a possible underlying genetic effect of adrenal reactivity to ACTH stimulation – but note that the sample size for the Franches-Montagnes horses was small at n=12, and 10 of these were driving horses trained by only three different trainers. Thus, results need to be interpreted with caution, and further confirmatory studies are needed.

Concerning the ‘number of riders’, interestingly, horses with two, three or more riders had a less pronounced adrenal response than horses with only one rider. Conceivably, horses that are accustomed to higher levels of stimulation in everyday life may be better equipped to deal with challenging situations [44–46]. In accordance, horses that spent more time outside of their stalls on pasture or for training and competition had a decreased reaction to exogenous ACTH. This is in accordance with previous studies indicating better welfare in pastured horses compared to stalled horses [6, 28, 47], possibly because the possibility to range outside more closely reflects the horse’s natural living conditions [6, 48].

‘Group housing’ was another variable that significantly influenced post-stimulatory cortisol concentrations. Horses living full-time in groups and horses living in groups on pasture showed higher post-stimulatory cortisol values than single housed horses. Contradictory results have been reported in previous studies investigating stress levels of domestic horses in different types of housing. One study found that horses living in single housing without physical contact to other horses had higher fecal glucocorticoids compared to horses with visual, auditory and tactile contact to other horses and horses living in groups of two or more individuals [22]. The authors concluded that integrating social contact into the housing of the horse improves equine welfare [22]. Other studies found no effect of the type of housing, neither in baseline salivary cortisol concentrations in adult horses [49] nor in fecal glucocorticoid metabolites and baseline salivary cortisol in young horses stabled for the first time [50, 51]. Similar to the results of the present study, Visser et al. [23] found lower cortisol concentrations in response to a CRH stimulation test in individually housed horses. In contrast to our interpretation, however, they proposed a decreased, not an increased response as a potential sign for chronic stress. In line with the analysis of the other variables in the present study, we conclude that group housing might represent a further stressor activating the HPA axis. Although group housing appears overall to have beneficial effects on horse welfare, it is also associated with social challenges [52], which might be reflected by the increased cortisol reactivity of group housed horses in the present study. In this context, social stability of groups might be important to consider in future studies, which itself may be influenced by many factors such as group size, age and sex composition, frequencies of changes or feeding management, available space and infrastructure.

Regarding personality, the present study could confirm the overall factor structure of the original questionnaire by Momozawa and colleagues [41] to some extent, although it yielded some additional factors and some differences in loadings of individual items. Some differences in factor structure are expectable given that firstly the study population was more varied than in Momozawa et al., where only racehorses were investigated, secondly not all questions were identical (two questions from the original version were removed and four questions added), and thirdly German and French translations of the questionnaire were used. Still, the similarity with the corresponding factors in Momozawa et al. indicates validity of the questionnaire. Nevertheless, the analysis revealed no significant associations between the cortisol response to ACTH and personality factors. Previous studies investigating associations between HPA axis function and personality traits in horses yielded diverse results. In line with our results, no correlations were found between temperament and basal serum cortisol values of horses used in therapeutic riding programs [53]. Another study investigated acute stress in stallions during transportation and found that calmer and more experienced stallions had a more pronounced increase in plasma cortisol following road transport than inexperienced, nervous stallions [54]. The authors concluded that lower cortisol concentrations in the latter group might represent a failure of the HPA axis to adapt to stress perceived during transportation [54]. As these examples demonstrate for horses, the evidence for associations between HPA axis function and personality has been inconclusive also in our own species [55] where both increased [33] and decreased cortisol concentrations [54] have been interpreted as signs of disturbed HPA function. Further research is needed in order to gain more specific insights into possible relations between HPA axis function and equine behavior.

The current study was the first study to compare stress of high-performance athletes with amateur sport horses and found no difference in post-stimulatory cortisol levels between the two groups. However, studies investigating acute stress in relation to training [56–58] and competition [29, 30] showed that experienced horses had lower basal plasma cortisol values than inexperienced horses [56]. Furthermore, horses with a superior physical training level had decreased basal cortisol levels, indicating that trained horses experience less stress during exercise than untrained horses [56–58]. Consequently, the increased requirements and the elevated stress levels that elite sport horses encounter may be compensated by their experience and their better fitness. This may be the reason why measurable indicators of long-term stress, such as cortisol responsiveness to external stimulation do not differ between elite and amateur sport horses. Overtraining, however, defined as a depletion of performance capacity, was revealed to lead to deleterious consequences including weight loss, behavioral changes and reluctance to exercise [59]. It was also reported to be associated with lower baseline cortisol values [36] and – quite contradictory – with an increased [60], decreased [61] or unchanged [36] adrenal response to exogenous ACTH. Interestingly, Bruin et al. [60] reported an increased response in horses with early stages of overtraining, whereas Persson et al. [61] found a decreased response in horses that were progressively and severely affected.

In comparison to these previous investigations, the present study tried to evaluate and compare the overall adrenal capacity of the sport horse in a normal environment without external stimuli such as training or transportation. Our results indicate that elite performance levels do not lead to increased adrenal cortex responsiveness to ACTH. We cannot specify, though, if this effect reflects targeted selection of horses that are able to withstand the enhanced requirements in high-performance competitions. Also, with respect to the effect of discipline, we found no difference in adrenal cortex reactivity between horses used for dressage, show jumping, eventing or endurance. While two previous studies reported increased cortisol levels in dressage horses [56, 62] compared to horses of other disciplines in a competition environment, our results indicate no long-term effect of discipline on HPA axis responsiveness.

A central limitation of this and similar studies remains the uncertain interpretation of the results of ACTH stimulation tests. While a decreased response has usually been interpreted as a sign of chronic stress in horses through desensitization of the HPA axis [23, 35, 61] (in its extreme form an Addison crisis), various studies demonstrate an increased response in potentially stressed horses [18–20, 22, 60]. Irrespective of these fundamental questions, a blunted response, typically described in the context of severe distress [63], was not expected in horses of the present study. At the time the study was conducted, all participating animals successfully took part in competitions and were in good health. Therefore, we hypothesized that increased responses might indicate higher stress responsiveness.

A further difficulty for the interpretation of the results is the use of an immunoassay for the measurement of cortisol concentrations in equine saliva. As shown for humans [64] and only recently for use in horses [65], cortisol results derived from immunoassays need to be interpreted with caution. While baseline values lack specificity, stimulated cortisol concentrations might be overestimated [65]. However, an evaluation of overall effects, at a group level rather than at an individual level, should still be reliable. Consequently, only stimulated cortisol concentrations were considered for the analysis of the present study. Previous studies described seasonal variability and a diurnal rhythm within cortisol concentrations [49, 66, 67]. Due to the need to adjust to horse owners’ and riders’ schedules, it was not possible to collect samples at the same time of the day or within only one season. However, baseline cortisol values, which are the most likely to be affected by natural occurring release patterns, were not analyzed here. Cortisol releases induced by exogenous ACTH administration are expected to override diurnal or seasonal changes [68], and we statistically controlled for this effect by including time of day as a predictor in the models. This confirmed that time of day had no significant effect on the post-stimulatory salivary cortisol levels reported here.

In conclusion, from the current data, there is no evidence that the superior performance and the high demands that elite sport horses have to fulfill in comparison to amateur sport horses have long-term adverse consequences on their welfare as determined by their cortisol reactivity. However, breed and management parameters, such as number of riders, time spent outside and housing, influence the amount of stress hormones released. Therefore, our results suggest that optimizing husbandry conditions, rather than changing performance level, may be more important for improving the welfare of horses.

## Acknowledgements

The authors thank the owners and riders of all participating horses for their help and commitment, as well the Swiss Equestrian Federation. In addition, they would like to thank the veterinarians Dr. Hervé Brünisholz, Dr. Michael Klopfenstein, Dr. Rolf Hegner and Dr. Selma Latif. By establishing contacts with participants, they were central to the success of this project. Great thanks also goes to Dr. Shannon Axiak Flammer, who helped to edit the manuscript, Lukas Bütikofer, PhD for his statistical advice and Prof. Rupert Bruckmaier and Chantal Phillipona, who enabled immunoassay sample analysis.

## References

1. Houpt KA. Domestic animal behavior for veterinarians and animal scientists: John Wiley & Sons; 2018.

2. Mayes E, Duncan P. Temporal Patterns of Feeding Behaviour in Free-Ranging Horses 1986. 105–29 p.

3. Hinchcliff KW, Kaneps AJ, Geor RJ. Equine sports medicine and surgery: basic and clinical sciences of the equine athlete: Elsevier Health Sciences; 2013.

4. Hinchcliff K, Geor R. The horse as an athlete: a physiological overview. Equine exercise physiology: the science of exercise in the athletic horse. 2008.

5. Art T, Lekeux P. Exercise-induced physiological adjustments to stressful conditions in sports horses. Livestock Production Science. 2005;92(2):101–11.

6. Werhahn H, Hessel EF, Van den Weghe HFA. Competition Horses Housed in Single Stalls (II): Effects of Free Exercise on the Behavior in the Stable, the Behavior during Training, and the Degree of Stress. Journal of Equine Veterinary Science. 2012;32(1):22–31.

7. Goodwin D, Davidson HP, Harris P. Foraging enrichment for stabled horses: effects on behaviour and selection. Equine Vet J. 2002;34(7):686–91.

8. McGowan C, Whitworth D. Overtraining syndrome in horses. Comparative Exercise Physiology. 2008;5(2):57–65.

9. Szabo S. Hans Selye and the development of the stress concept. Special reference to gastroduodenal ulcerogenesis. Ann N Y Acad Sci. 1998;851(1):19–27.

10. Borstel UKv, Visser E, Hall C. Indicators of stress in equitation. Applied Animal Behaviour Science. 2017;190:43–56.

11. Moberg GP. Biological response to stress: implications for animal welfare. The biology of animal stress: basic principles and implications for animal welfare. 2000:1–21.

12. Bachmann I, Bernasconi P, Herrmann R, Weishaupt M, Stauffacher M. Behavioural and physiological responses to an acute stressor in crib-biting and control horses. Applied Animal Behaviour Science. 2003;82(4):297–311.

13. Malmkvist J, Poulsen JM, Luthersson N, Palme R, Christensen JW, Sondergaard E. Behaviour and stress responses in horses with gastric ulceration. Applied Animal Behaviour Science. 2012;142(3-4):160–7.

14. Schmidt A, Aurich J, Mostl E, Muller J, Aurich C. Changes in cortisol release and heart rate and heart rate variability during the initial training of 3-year-old sport horses. Horm Behav. 2010;58(4):628–36.

15. Ladewig J, Smidt D. Behavior, episodic secretion of cortisol, and adrenocortical reactivity in bulls subjected to tethering. Horm Behav. 1989;23(3):344–60.

16. Van Reenen C, Mars M, Leushuis I, Rijsewijk F, Van Oirschot J, Blokhuis H. Social isolation may influence responsiveness to infection with bovine herpesvirus 1 in veal calves. Veterinary microbiology. 2000;75(2):135–43.

17. Hedberg Y, Dalin AM, Forsberg M, Lundeheim N, Hoffmann B, Ludwig C, et al. Effect of ACTH (tetracosactide) on steroid hormone levels in the mare. Part A: effect in intact normal mares and mares with possible estrous related behavioral abnormalities. Anim Reprod Sci. 2007;100(1-2):73–91.

18. Freymond SB, Bardou D, Briefer EF, Bruckmaier R, Fouche N, Fleury J, et al. The physiological consequences of crib-biting in horses in response to an ACTH challenge test. Physiology & Behavior. 2015;151:121–8.

19. Sauer FJ, Bruckmaier RM, Ramseyer A, Vidondo B, Scheidegger MD, Gerber V. Diagnostic accuracy of post-ACTH challenge salivary cortisol concentrations for identifying horses with equine glandular gastric disease. Journal of animal science. 2018;96(6):2154–61.

20. Scheidegger MD, Gerber V, Bruckmaier RM, van der Kolk JH, Burger D, Ramseyer A. Increased adrenocortical response to adrenocorticotropic hormone (ACTH) in sport horses with equine glandular gastric disease (EGGD). Vet J. 2017;228:7–12.

21. Von Borell E, Ladewig J. Altered adrenocortical response to acute stressors or ACTH (1-24) in intensively housed pigs. Domestic animal endocrinology. 1989;6(4):299–309.

22. Yarnell K, Hall C, Royle C, Walker SL. Domesticated horses differ in their behavioural and physiological responses to isolated and group housing. Physiology & Behavior. 2015;143:51–7.

23. Visser EK, Ellis AD, Van Reenen CG. The effect of two different housing conditions on the welfare of young horses stabled for the first time. Applied Animal Behaviour Science. 2008;114(3-4):521–33.

24. McGreevy PD, Cripps PJ, French NP, Green LE, Nicol CJ. Management factors associated with stereotypic and redirected behaviour in the thoroughbred horse. Equine Vet J. 1995;27(2):86–91.

25. Parker M, Goodwin D, Redhead ES. Survey of breeders’ management of horses in Europe, North America and Australia: comparison of factors associated with the development of abnormal behaviour. Applied Animal Behaviour Science. 2008;114(1-2):206–15.

26. Bachmann I, Audige L, Stauffacher M. Risk factors associated with behavioural disorders of crib-biting, weaving and box-walking in Swiss horses. Equine Vet J. 2003;35(2):158–63.

27. Christie JL, Hewson CJ, Riley CB, McNiven MA, Dohoo IR, Bate LA. Management factors affecting stereotypies and body condition score in nonracing horses in Prince Edward Island. Can Vet J. 2006;47(2):136–43.

28. McGreevy PD, French NP, Nicol CJ. The prevalence of abnormal behaviours in dressage, eventing and endurance horses in relation to stabling. Vet Rec. 1995;137(2):36–7.

29. Becker-Birck M, Schmidt A, Lasarzik J, Aurich J, Mostl E, Aurich C. Cortisol release and heart rate variability in sport horses participating in equestrian competitions. Journal of Veterinary Behavior-Clinical Applications and Research. 2013;8(2):87–94.

30. Janczarek I, Bereznowski A, Strzelec K. The influence of selected factors and sport results of endurance horses on their saliva cortisol concentration. Polish Journal of Veterinary Sciences. 2013;16(3):533–41.

31. Bartolome E, Cockram MS. Potential Effects of Stress on the Performance of Sport Horses. Journal of Equine Veterinary Science. 2016;40:84–93.

32. Valenchon M, Levy F, Moussu C, Lansade L. Stress affects instrumental learning based on positive or negative reinforcement in interaction with personality in domestic horses. PLoS One. 2017;12(5):e0170783.

33. Valenchon M, Levy F, Prunier A, Moussu C, Calandreau L, Lansade L. Stress modulates instrumental learning performances in horses (Equus caballus) in interaction with temperament. PLoS One. 2013;8(4):e62324.

34. Valenchon M, Levy F, Fortin M, Leterrier C, Lansade L. Stress and temperament affect working memory performance for disappearing food in horses, Equus caballus. Animal Behaviour. 2013;86(6):1233–40.

35. Hoffman RM, Kronfeld DS, Holland JL, Greiwe-Crandell KM. Preweaning diet and stall weaning method influences on stress response in foals. J Anim Sci. 1995;73(10):2922–30.

36. Golland LC, Evans DL, Stone GM, Tyler-McGowan CM, Hodgson DR, Rose RJ. Plasma cortisol and ß-endorphin concentrations in trained and over-trained Standardbred racehorses. Pflügers Archiv European Journal of Physiology. 1999;439(1):11–7.

37. Wagner EL, Tyler PJ. A Comparison of Weight Estimation Methods in Adult Horses. Journal of Equine Veterinary Science. 2011;31(12):706–10.

38. Carroll CL, Huntington PJ. Body condition scoring and weight estimation of horses. Equine Vet J. 1988;20(1):41–5.

39. Peeters M, Sulon J, Beckers JF, Ledoux D, Vandenheede M. Comparison between blood serum and salivary cortisol concentrations in horses using an adrenocorticotropic hormone challenge. Equine Veterinary Journal. 2011;43(4):487–93.

40. Scheidegger MD, Gerber V, Ramseyer A, Schupbach-Regula G, Bruckmaier RM, van der Kolk JH. Repeatability of the ACTH stimulation test as reflected by salivary cortisol response in healthy horses. Domestic Animal Endocrinology. 2016;57:43–7.

41. Momozawa Y, Kusunose R, Kikusui T, Takeuchi Y, Mori Y. Assessment of equine temperament questionnaire by comparing factor structure between two separate surveys. Applied Animal Behaviour Science. 2005;92(1-2):77–84.

42. R Core Team. R: A Language and Environment for Statistical Computing. R Foundation for Statistical Computing. 2019.

43. Sauer FJ, Bruckmaier RM, Ramseyer A, Vidondo B, Scheidegger MD, Gerber V. Diagnostic accuracy of post-ACTH challenge salivary cortisol concentrations for identifying horses with equine glandular gastric disease. J Anim Sci. 2018;96(6):2154–61.

44. Fox C, Merali Z, Harrison C. Therapeutic and protective effect of environmental enrichment against psychogenic and neurogenic stress. Behavioural brain research. 2006;175(1):1–8.

45. Parker KJ, Buckmaster CL, Schatzberg AF, Lyons DM. Prospective investigation of stress inoculation in young monkeys. Archives of general psychiatry. 2004;61(9):933–41.

46. Brockhurst J, Cheleuitte-Nieves C, Buckmaster C, Schatzberg A, Lyons D. Stress inoculation modeled in mice. Translational psychiatry. 2015;5(3):e537.

47. Rivera E, Benjamin S, Nielsen B, Shelle J, Zanella AJ. Behavioral and physiological responses of horses to initial training: the comparison between pastured versus stalled horses. Applied Animal Behaviour Science. 2002;78(2-4):235–52.

48. Kiley-Worthington M. The behavior of horses in relation to management and training— towards ethologically sound environments. Journal of Equine Veterinary Science. 1990;10(1):62–75.

49. Aurich J, Wulf M, Ille N, Erber R, von Lewinski M, Palme R, et al. Effects of season, age, sex, and housing on salivary cortisol concentrations in horses. Domest Anim Endocrinol. 2015;52:11–6.

50. Heleski CR, Shelle AC, Nielsen BD, Zanella AJ. Influence of housing on weanling horse behavior and subsequent welfare. Applied Animal Behaviour Science. 2002;78(2-4):291–302.

51. Harewood EJ, McGowan CM. Behavioral and physiological responses to stabling in naive horses. Journal of Equine Veterinary Science. 2005;25(4):164–70.

52. Alexander SL, Irvine CH. The effect of social stress on adrenal axis activity in horses: the importance of monitoring corticosteroid-binding globulin capacity. J Endocrinol. 1998;157(3):425–32.

53. Anderson MK, Friend TH, Evans JW, Bushong DM. Behavioral assessment of horses in therapeutic riding programs. Applied Animal Behaviour Science. 1999;63(1):11–24.

54. Fazio E, Medica P, Cravana C, Ferlazzo A. Cortisol response to road transport stress in calm and nervous stallions. Journal of Veterinary Behavior-Clinical Applications and Research. 2013;8(4):231–7.

55. Hauner KK, Adam EK, Mineka S, Doane LD, DeSantis AS, Zinbarg R, et al. Neuroticism and introversion are associated with salivary cortisol patterns in adolescents. Psychoneuroendocrinology. 2008;33(10):1344–56.

56. Cayado P, Munoz-Escassi B, Dominguez C, Manley W, Olabarri B, Sanchez de la Muela M, et al. Hormone response to training and competition in athletic horses. Equine Vet J Suppl. 2006;38(36):274–8.

57. Marc M, Parvizi N, Ellendorff F, Kallweit E, Elsaesser F. Plasma cortisol and ACTH concentrations in the warmblood horse in response to a standardized treadmill exercise test as physiological markers for evaluation of training status. Journal of Animal Science. 2000;78(7):1936–46.

58. Mircean M, Giurgiu G, Mircean V, Zinveliu E. Serum cortisol variation of sport horses in relation with the level of training and effort intensity. Bulletin USAMV-CN. 2007;64(1-2):488–92.

59. Castejon-Riber C, Riber C, Rubio MD, Aguera E, Munoz A. Objectives, Principles, and Methods of Strength Training for Horses. Journal of Equine Veterinary Science. 2017;56:93–103.

60. Bruin G, Kuipers H, Keizer HA, Vander Vusse GJ. Adaptation and overtraining in horses subjected to increasing training loads. J Appl Physiol (1985). 1994;76(5):1908–13.

61. Persson SG, Larsson M, Lindholm A. Effects of training on adreno-cortical function and red-cell volume in trotters. Zentralbl Veterinarmed A. 1980;27(4):261–8.

62. Munk R, Jensen RB, Palme R, Munksgaard L, Christensen JW. An exploratory study of competition scores and salivary cortisol concentrations in Warmblood horses. Domest Anim Endocrinol. 2017;61:108–16.

63. Pawluski J, Jego P, Henry S, Bruchet A, Palme R, Coste C, et al. Low plasma cortisol and fecal cortisol metabolite measures as indicators of compromised welfare in domestic horses (Equus caballus). Plos One. 2017;12(9):e0182257.

64. Bae YJ, Gaudl A, Jaeger S, Stadelmann S, Hiemisch A, Kiess W, et al. Immunoassay or LC-MS/MS for the measurement of salivary cortisol in children? Clin Chem Lab Med. 2016;54(5):811–22.

65. Sauer FJ, Gerber V, Frei S, Bruckmaier RM, Groessl M. Salivary cortisol measurement in horses: Immunoassay or LC-MS/MS? submitted.

66. Cordero M, Brorsen BW, McFarlane D. Circadian and circannual rhythms of cortisol, ACTH, and alpha-melanocyte-stimulating hormone in healthy horses. Domest Anim Endocrinol. 2012;43(4):317–24.

67. Irvine CH, Alexander SL. Factors affecting the circadian rhythm in plasma cortisol concentrations in the horse. Domest Anim Endocrinol. 1994;11(2):227–38.

68. Schmidt A, Aurich C, Neuhauser S, Aurich J, Möstl E, editors. Comparison of cortisol levels in blood plasma, saliva and faeces of horses submitted to different stressors or treated with ACTH. Proceedings, 5 Intern Symposium Equitation Science; 2009; Sydney.

